## Supplementary figures and images for "Cellular and molecular characterization of peripheral glia in the lung and other organs"

### SuppFiles_combined

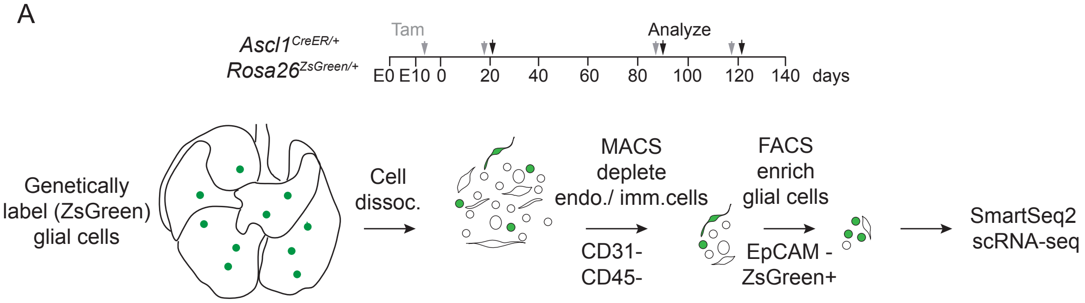

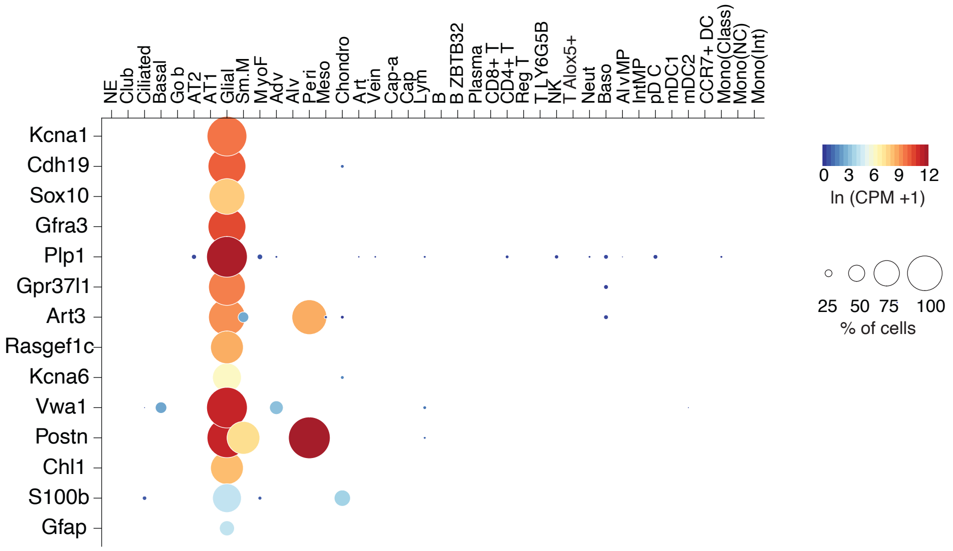

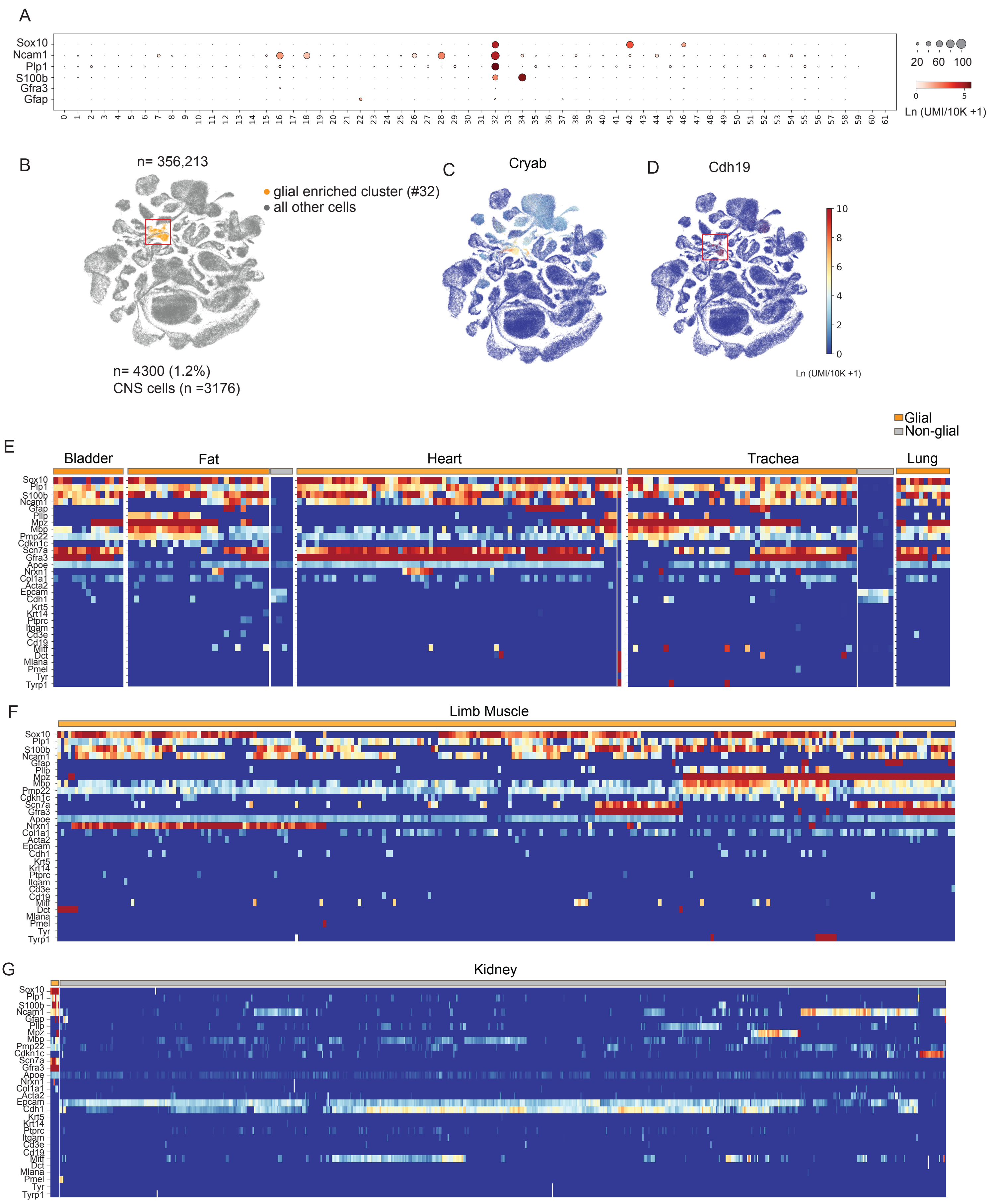

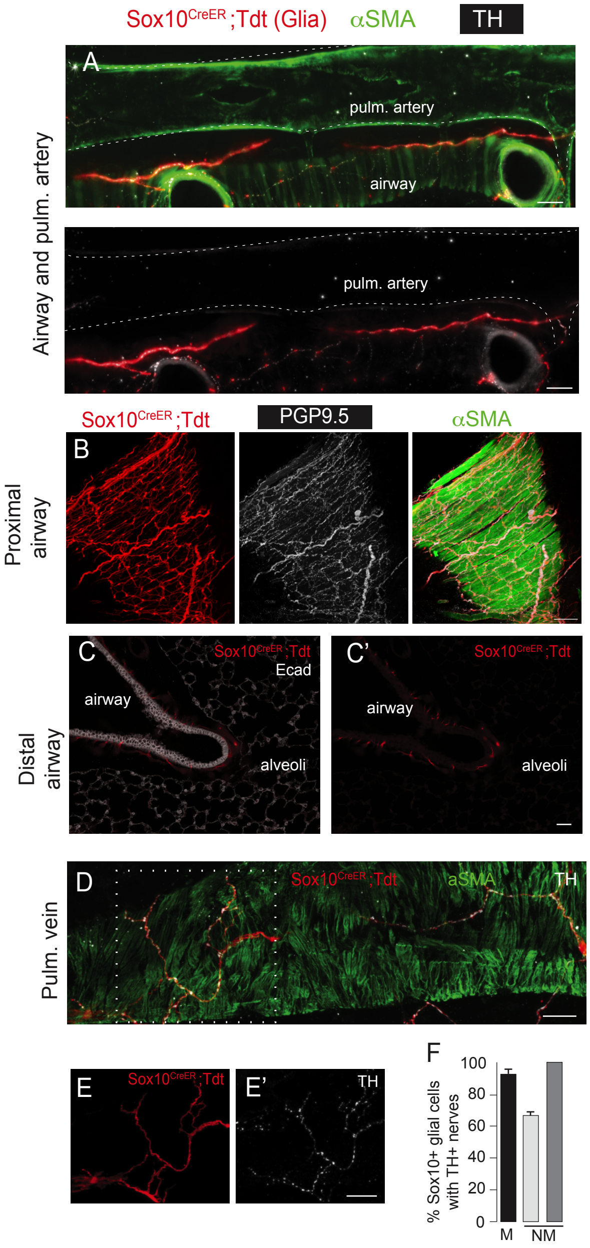

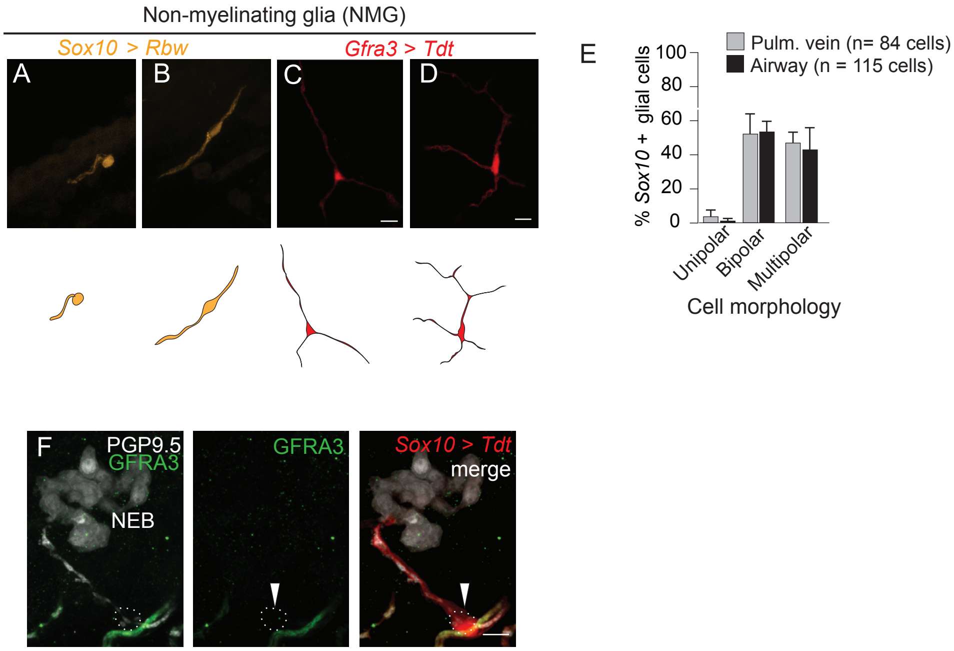

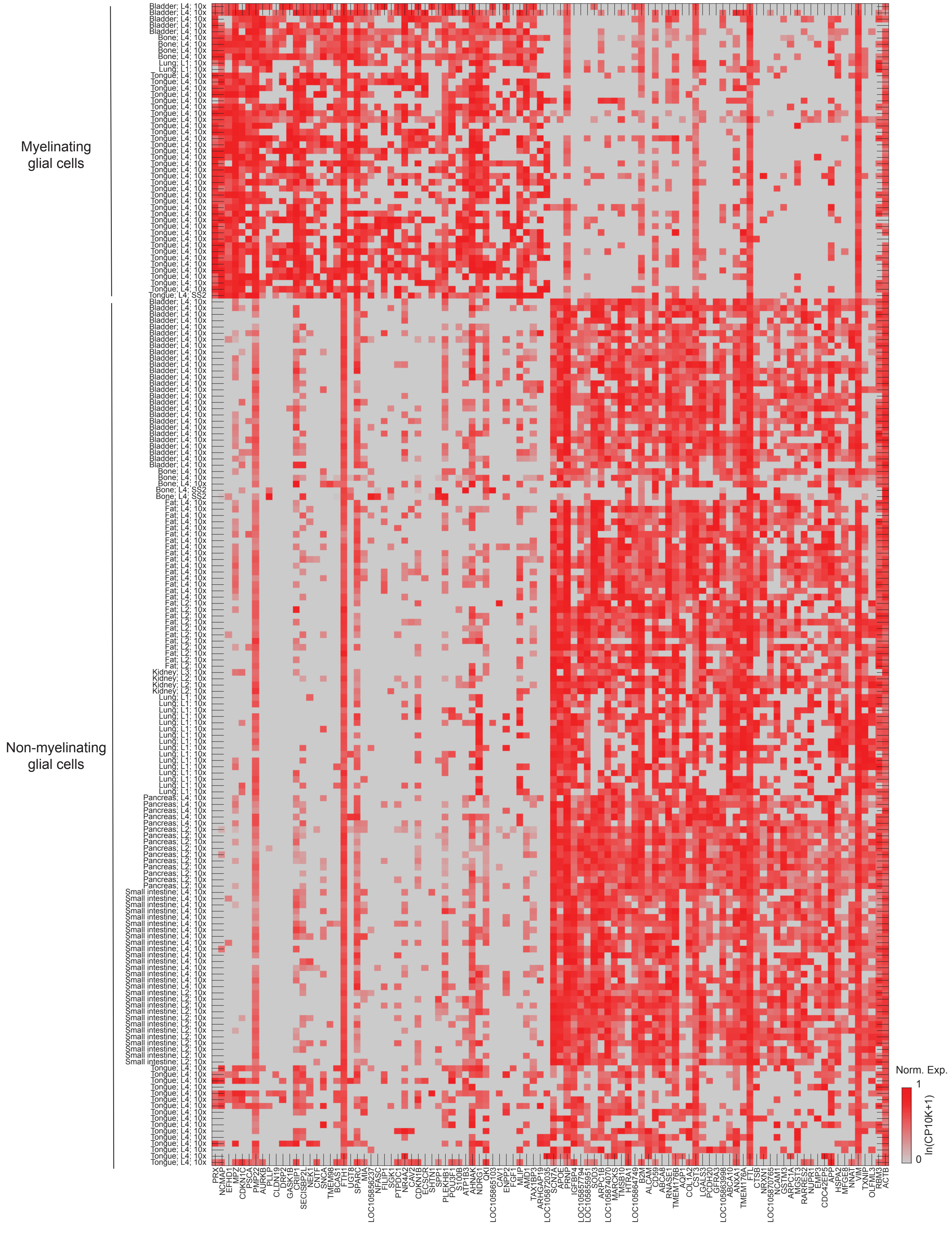

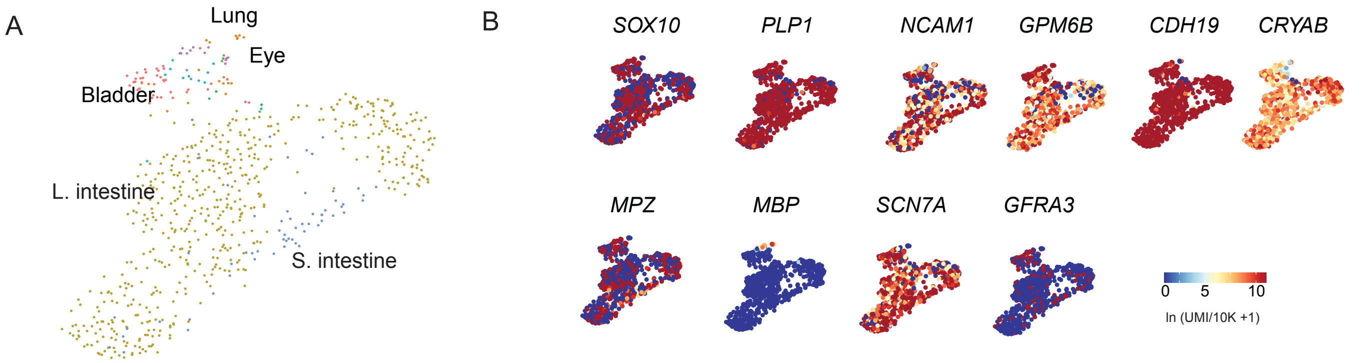

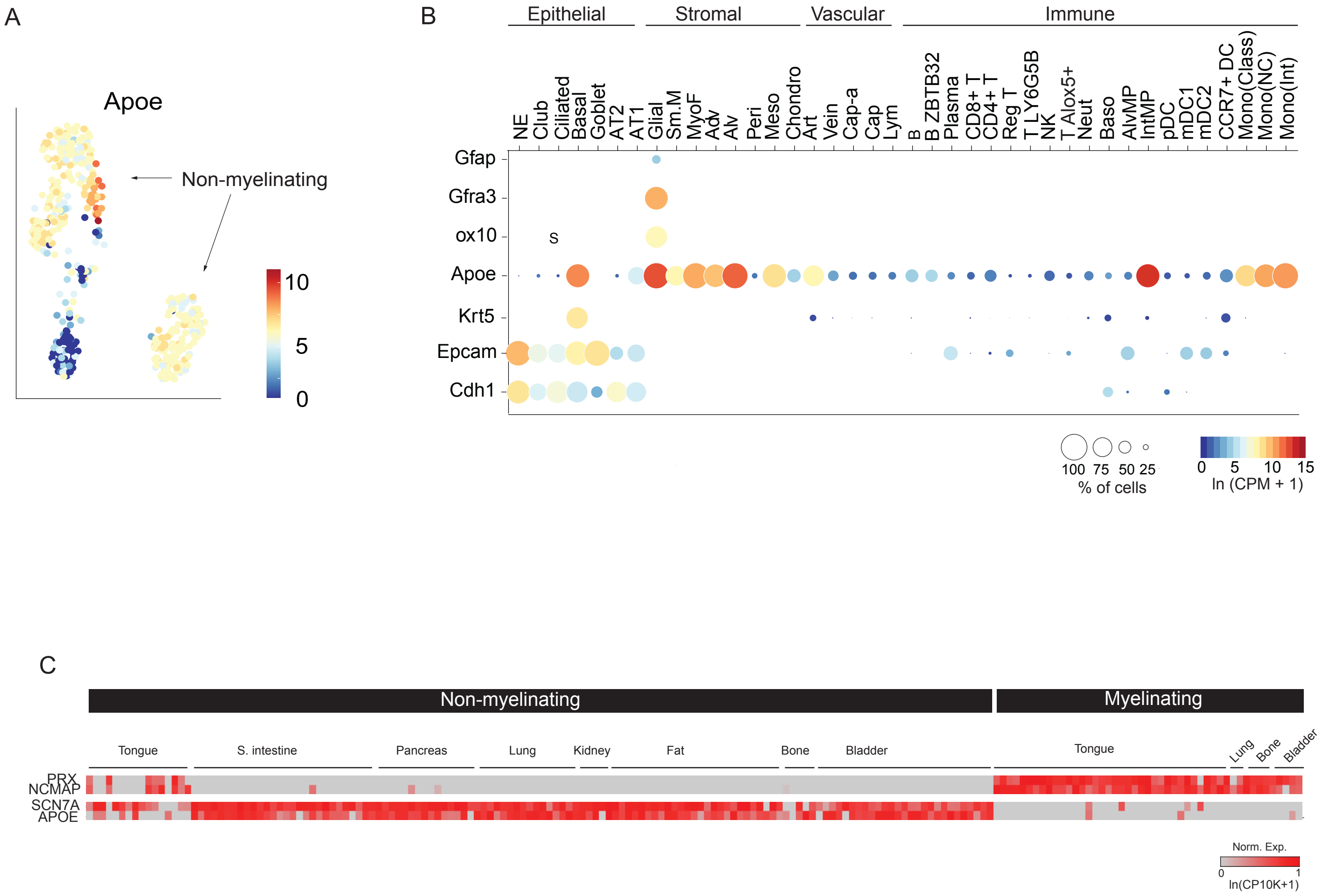

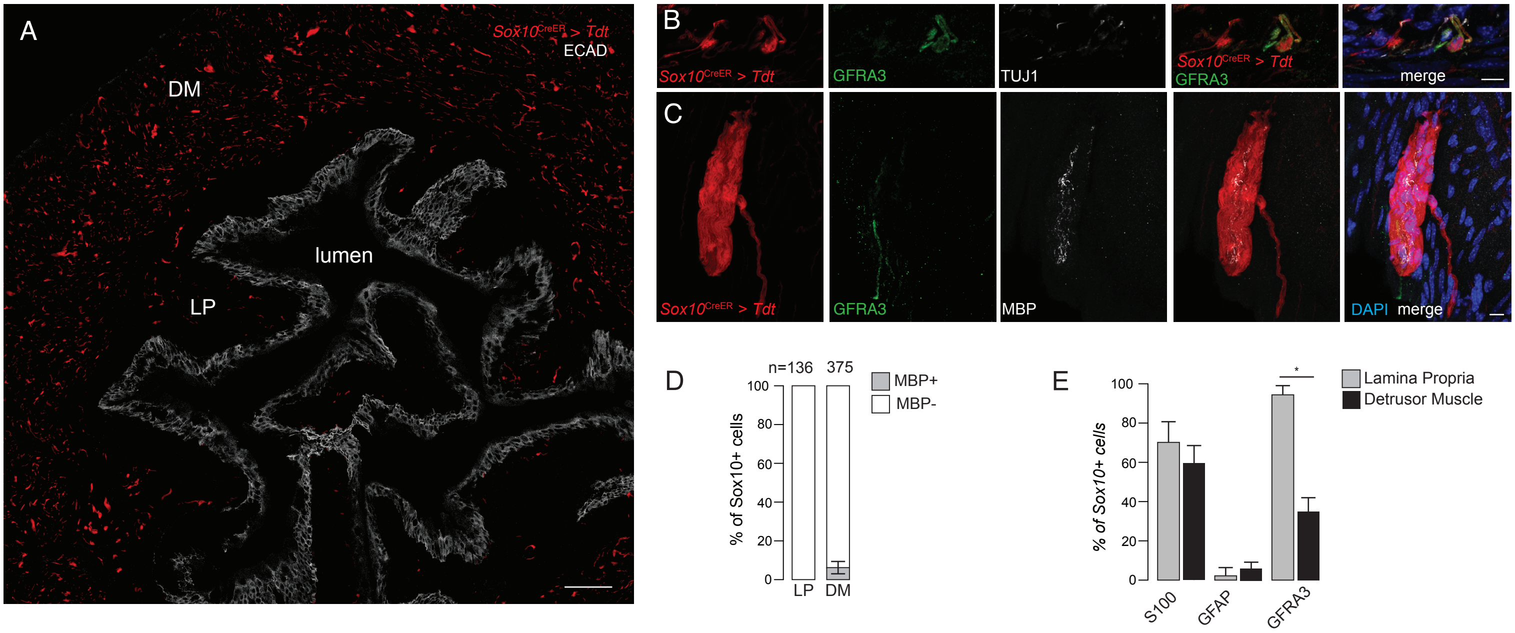
